# Humic substances mitigate adverse effects of elevated temperature with potentially critical repercussions for coral reef resilience

**DOI:** 10.1101/2023.04.14.536861

**Authors:** T.M. Stuij, D.F.R. Cleary, N.J. de Voogd, R.J.M. Rocha, A.R.M. Polonia, D.A.M. Silva, J.C. Frommlet, A. Louvado, Y. M. Huang, N. van der Windt, N.C.M. Gomes

**Affiliations:** Centre for Environmental and Marine Studies (CESAM) and Department of Biology, University of Aveiro, Aveiro, Portugal; Naturalis Biodiversity Center, Leiden, The Netherlands; Institute of Environmental Sciences (CML), Leiden University, Leiden, The Netherlands; Penghu University of Science and Technology, Magong, Taiwan

**Keywords:** bacterial communities, microbes, terrestrial organic matter, climate change, ENSO, coastal tropical coral reefs, *Montipora digitata*, *Montipora capricornis*, *Sarcophyton glaucum*, *Chondrilla* sp

## Abstract

Previous observational studies have suggested that terrestrially-derived compounds, most notably humic substances (HS) can protect coral reefs from thermal stress. No study hitherto has, however, tested this hypothesis. In the present study, we used a randomised-controlled microcosm setup to test to what extent HS are able to mitigate the adverse effects of elevated temperature and intense UVB radiation on coral photosynthetic activity, and environmental and host-associated bacterial ercommunities. Our results clearly demonstrate a significant protective effect of HS. Corals in HS-supplemented microcosms had significantly higher photosynthetic activities than those in microcosms subjected to elevated heat and intense UVB radiation. Our results, furthermore, showed that coral reef organisms in HS-supplemented microcosms contained unique bacterial communities enriched with known groups of potentially beneficial bacteria. Our findings have significant repercussions for reef resilience in the face of increasing climate-induced stressors and highlight the importance of restoring coastal forests and the land-sea interface in order to protect coral reefs.

## 1. Introduction

Coral reefs are one of the most diverse of marine ecosystems and provide numerous ecological and economic services to adjacent communities (Moberg and Folke, 1999). The intricate calcium carbonate coral skeletons provide the building blocks for highly complex three-dimensional reef structures, and habitat for numerous species (Roberts et al., 2002). Many corals form obligate symbioses with dinoflagellates, also known as zooxanthellae (Trench, 1979). These unicellular algae photosynthesize and provide sugars and other nutrients to their hosts (Jones & Yellowlees, 1997). Zooxanthellae are of such high importance to their hosts, that disturbances to the symbiotic relationship can lead to host demise (Glynn, 1984). One of the best-known examples of a breakdown of symbiosis is coral bleaching. Large-scale bleaching events are tightly linked to elevated temperatures and intense solar radiation, particularly in relation to extreme El Niño Southern Oscillation (ENSO) events (Hughes et al., 2017). As the frequency and severity of such events is predicted to increase in coming decades (Ying et al., 2022), the future state of coral reef ecosystems will be determined by how they respond to these environmental disturbances.

In addition to zooxanthellae, bacterial symbionts inhabiting coral tissue have been shown to play key roles in coral response to environmental perturbations (Doering et al., 2021; Rosado et al., 2019). For example, the degree of bleaching of the coral *Poccilopora damicornis* was found to be associated with the presence and/or abundances of several bacterial taxa, such as *Pseudoalteromonas* spp., *Halomonas taeanensis* and *Colbetia marina* (Rosado et al., 2019). Moreover, bacterial symbionts play key roles in important processes including nitrogen fixation and sulfur cycling, and provide their coral hosts with essential vitamins and antioxidants; they also play roles in pathogen defense and coral health in general (Glasl et al., 2016). In addition to bacterial symbionts, coral-associated bacterial communities may also include opportunistic pathogens. Members of the families *Vibrionaceae* and *Alteromonadaceae*, for example, have repeatedly been shown to colonize the tissues of corals exposed to unfavorable conditions (e.g., increasing water temperatures) (Bourne et al., 2016; McDevitt-Irwin et al., 2017). In addition to corals, bacterial symbionts have also been shown to play key roles in the health of a wide range of other hosts from sponges to humans (Gilbert et al., 2018; Posadas et al., 2022).

Recent studies have attempted to manipulate coral microbiomes in order to enhance coral resistance and resilience to severe environmental stressors and pathogens (Doering et al., 2021; Rosado et al., 2019; Ziegler et al., 2019). However, the use of microbiome modulation approaches based on the use of living microorganisms in coral reefs is challenging due to the potential unintended effects of introducing putative probiotics to other cohabiting organisms and the lack of established methodologies for effective introduction of these microorganisms (Sweet et al., 2017). Chemical microbiome modulators, for example prebiotics, that promote the growth of naturally occurring beneficial microbes, can be a more natural and practical approach to stimulate the growth of beneficial bacterial taxa. HS, for example, have been previously used to improve soil properties, plant growth and, importantly, soil bacterial diversity in agricultural systems (da Silva et al., 2021; Nardi et al., 2021). They have, however, received limited attention as modulators of bacterial communities in aquatic systems (Louvado et al., 2021).

HS are complex organic compounds that are formed through the decomposition of plant organic matter in terrestrial ecosystems (MacCarthy, 2001). Terrestrial-derived HS mainly enter the marine environment via underwater cave systems or river runoff, and are found in greater concentrations near coastal ecosystems (Esham et al., 2000). Concentrations decline with increasing distance to land (Esham et al., 2000).

HS influence the photosynthetically active radiation (PAR; 400 – 700 nm) in the water column and benthic substrate, which subsequently affects photoautotrophs, such as phytoplankton communities and algal coral symbionts (Sharpless et al., 2014; Gerea et al., 2017; Roth 2014). In addition to this, HS absorb sunlight in the ultraviolet (UV) spectrum, particularly UVB wavelengths (280 – 320 nm), which is attributed to aromatic chromophores (Sharpless et al., 2014), and is thought to convey UV-protective properties (Ayoub et al., 2012; errier-Pagès et al., 2007).

HS have also been shown to affect marine bacterioplankton community composition (Lindh et al., 2015) and to favor the growth of particular groups of potentially beneficial bacteria belonging to the Alphaproteobacteria (e.g., the *Roseobacter* clade and candidate divisions SAR11 and SAR86) in addition to others (Esham et al., 2000; Lindh et al., 2015; Rocker et al., 2012). Recently, Louvado et al. (2021) showed that HS modulated fish skin mucus bacterial communities and increased the relative abundances of putative beneficial bacteria belonging to the *Roseobacter* clade. However, to the best of our knowledge, no studies have been conducted to assess the effects of HS on environmental and host-associated bacterial communities in a coral reef setting.

Marine ecosystems have been historically affected by destruction of coastal and riparian ecosystems (Stoms et al., 2005). Coastal forests, mangroves and seagrass meadows have been subjected to large-scale degradation and loss of habitat in order to provide space for urban development, industries and intensive agri- and aquaculture (Crain et al., 2009). This development has resulted in increased transfer of nutrients and pollutants to coastal marine environments (Petersen et al., 2017), adversely affecting corals that would normally thrive under oligotrophic conditions (Burkepile et al., 2020; Ferrier-Pagès et al., 2000; Hall et al., 2018). This development has, furthermore, resulted in qualitative and quantitative shifts in HS that are transferred to the adjacent marine environment (dos Santos et al., 2019; Spaccini et al., 2006). For example, soils of deforested agricultural fields showed a 38 to 53% reduction in HS relative to soils of natural forests (dos Santos et al., 2019). Although the impact of nutrient enrichment on tropical coral reefs has been studied extensively (Burkepile et al., 2020; Ferrier-Pagès et al., 2000; Hall et al., 2018), the specific effects related to the loss and modification of HS in these ecosystems are relatively unknown. Observational studies (Ayoub et al. 2012; Cleary et al. 2016) have, however, suggested a protective effect of HS in the form of chromophoric dissolved organic matter, also known as CDOM, a riverine input tracer (Fichot et al., 2013). Hitherto, however, no study has explicitly tested for such a protective effect. High CDOM concentrations can result from decaying plant material transported via land run-off from areas with high vegetation productivity or from mangroves and seagrasses (Richardson and LeDrew, 2006). Importantly, Cleary et al. (2016) discussed the elevated CDOM levels in offshore coral reefs adjacent to the city of Jakarta, Indonesia and that remotely sensed images indicated that they derived from plumes emanating from the still, largely forested Borneo as opposed to the denuded coastline of Java. Coral reefs had already largely disappeared from inshore areas of Jakarta Bay prior to 1985, although the area once hosted vibrant reefs (Cleary et al. 2014).

The present study is the first to test if HS are able to mitigate the adverse effects of elevated temperature and intense UVB radiation on a key indicator of coral health, namely, photosynthetic activity in addition to assessing the independent and combined effects of temperature, UVB radiation and HS supplementation on environmental and host-associated bacterial communities. The organisms studied included two stony coral species, one soft coral and one sponge species and represent a range of taxa commonly found in Indo-pacific coral reef environments. Our results highlight a potential link between large-scale clearance of coastal/riparian forests and changes in HS-mediated effects on coral reef microbiomes and resilience.

With the present study, we tested the following hypotheses 1. HS will mitigate the effects of elevated temperature and UVB radiation on coral photosynthetic activity. 2. Elevated temperature and UVB radiation will induce significant shifts in the composition of host-associated and environmental (sediment and seawater) bacterial communities. 3. HS will mitigate the effects of elevated temperature and UVB radiation on environmental and host-associated bacterial communities whereby communities subjected to HS supplementation will more closely resemble communities kept under natural conditions (e.g., normal temperature).

## 2. Methods

### 2.1 ELSS design

The ELSS developed in this study was based on a microcosm system previously designed to assess the effects of global climate change and environmental contamination on sediment communities (Coelho et al., 2013). This system was modified and validated for coral reef microcosm experiments under laboratory-controlled conditions as described in detail in Stuij et al. (2023). Briefly, the ELSS included 32 glass aquaria (referred to as microcosms, 23 cm in height, 16 cm length and 12 cm width), which were individually connected to another aquarium (referred to as reservoir, 30 cm in height, 12 cm length and 12 cm width). Each reservoir-microcosm unit contained a functional water volume of ∼5 L. Water was circulated within microcosm-reservoir units using small hydraulic pumps, resulting in a constant flow rate of 8.64 ml/s (s = 0.66 ml/s). Water temperature was regulated using water bath tanks each surrounding four microcosm-reservoir units, equipped with water heaters with an internal thermostat (V2Therm 100 Digital heater). Constant aeration within microcosms was maintained using diffusing air stones connected to an air pump (530L/h, Resun) via small hoses. Lighting was controlled by four programmable luminaire systems (Reef - SET, Rees, Germany), each holding eight fluorescent lamps (Coelho et al., 2013). During the current experiment, four UV fluorescent tubes (SolarRaptor, T5/54W, Rees, Germany) and four full spectra fluorescent tubes (ATI AquaBlue Special, T5/54W) were connected alternately and programmed to a 12 h diurnal light cycle, simulating photoperiod conditions of tropical latitudes. To block radiation in the UVB spectrum (290 – 320 nm), microcosms were covered independently from each other by a transparent polyester film (Folanorm SF-AS, Folex coating, Köln, Germany). The film has been previously used in multiple studies to block UVB radiation (Müller et al., 2009; Rautenberger et al., 2013). In our experimental set-up, the film absorbed 90% of the UVB, 31% of UVA and 9% of PAR irradiance.

Using the ELSS, a multi-factorial experiment was designed for the testing of independent and interactive effects of HS supplementation, temperature and UVB radiation. Each factor had two levels, namely, HS supplementation (with *versus* without), temperature (28 °C *versus* 32 °C) and UVB radiation (with *versus* without) for a total of eight treatments with four replicates each. The full experiment, thus, consisted of 32 microcosms. The temperature treatment was randomized in groups of four microcosms, whereby HS supplementation, UVB radiation and the combination of both were randomly assigned within each group of microcosms with equal temperature. A graphical summary of the ELSS and different phases of the experiment can be found in Fig. 1.

**Fig. 1.**
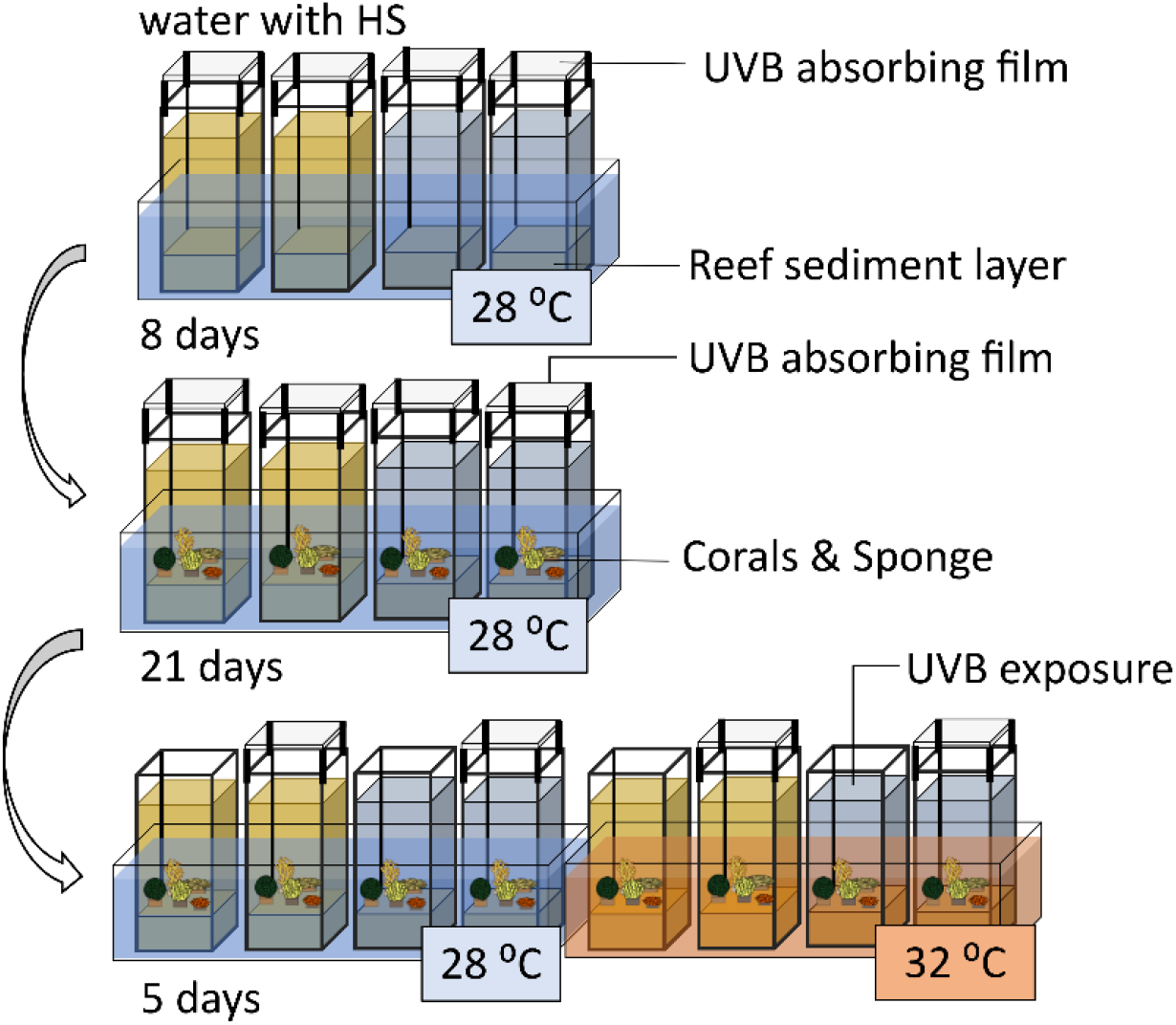
Graphical representation of the experimental set-up.

Each microcosm received a coral reef sediment layer of ∼3 cm, consisting of a mixture of commercially available (Reef Pink dry aragonite sand, Red Sea) and natural coral reef sediment. The commercial sediment was washed and sterilized (3 times autoclavation at 121ºC for 20 minutes (Otte et al., 2018) before use. The natural sediment was collected from a coral reef south of Fongguei, Penghu, Taiwan (22°19’50.5”N 120°22’19.8”E). Synthetic seawater used in the experiment was prepared by mixing coral reef salt (CORAL PRO SALT, Red Sea) with deionized water (V2Pure 360), with salinity adjusted to 35 ppt. A concentrated HS stock solution (10 gL^-1^) was first prepared by dissolving commercially available humic substances (technical grade humic acid, Sigma-Aldrich) in 0.2M NaOH water solution. Subsequently, the solution was neutralized to a pH of 8.0 – 8.2 by adding concentrated HCL. This concentrated stock solution was then used to enrich the synthetic seawater to a final HS concentration of 7.5 mgL^-1^. This concentration falls within the range of dissolved organic carbon (DOC) concentrations measured in coastal coral reef ecosystems, which were shown to vary from 0.05 to up to 17 mgL^-^1 (Kegler et al., 2018; Teichberg et al., 2018). Synthetic seawater with and without HS was poured into microcosms containing sediment, and the temperature was set to 28 °C. Partial seawater renewal was carried out daily by replacing 1 L of seawater with newly prepared synthetic seawater, with or without HS, in each microcosm. HS concentration within the water column of the microcosms was monitored using UV spectrometry (Eaton, 1995). The absorbance at a wavelength of 300 nm was relatively stable over the course of the experiment and averaged 0.017 ± 0.003 AU among microcosms (referenced against non HS-supplemented microcosm water).

After eight days of initial stabilization of the system, we added five reef organisms to each individual microcosm. The species were previously grown on carbonate stones (3 cm in diameter, 1 cm thickness) and included two hard corals, *Montipora digitata* (Dana, 1846) and *Montipora capricornis* Veron 1985, one soft coral, *Sarcophyton glaucum* (Quoy & Gaimard, 1833), one zoanthid, *Zoanthus* sp., and one sponge, *Chondrilla* sp. (Schmidt, 1862). All animals used in this study were obtained from the collection of marine invertebrates cultivated at ECOMARE (University of Aveiro, Portugal). ECOMARE holds validated coral reef culture systems (Rocha et al., 2015), which have previously been used to study environmental effects on coral reef species under experimentally controlled conditions (Rocha et al., 2020).

After adding the animals, the microcosms were kept in the conditions mentioned above for an additional 21 days. After this period, temperature and UVB treatments were applied to the microcosms for five days as follows. In the heat-treated microcosms, temperature was gradually increased over the course of three days from 28.0 ± 0.6 °C at day 1 to 31.0 ± 0.6 °C at day 3 and averaged 31.4 ± 0.5 °C and 31.2 ± 0.7 °C at days 4 and 5. These temperatures were chosen to match temperatures previously observed in Pacific coral reefs, which ranged between 26 °C to 32 °C (Kusuma et al., 2017). In the UVB treated microcosms, the UVB absorbing film was removed, which resulted in daily doses of 2.43 * 10 ^-2^ Jcm^-2^ UVB radiation (fig. 1).

### 2.2. In vivo chlorophyll fluorescence analysis

At the end of the experiment, chlorophyll fluorescence of the corals was measured in vivo using a pulse amplitude modulation (PAM) fluorometer (Walz™). Fluorescence was measured in dark-adapted samples (for 20 min), with Junior PAM and WinControl3 software (Walz™). Saturating light pulses (450 nm) were performed perpendicularly to the sample surface, with a 1.5 mm fiber optic. The maximum quantum yield (Fv/Fm) of photosystem II was calculated as 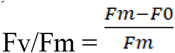.

### 2.3. Bacterial community analysis

#### 2.3.1. Sampling and DNA extraction

Samples of the sediment, water and organisms were collected five days after the beginning of the experiment. A composite sediment sample was collected from each microcosm by haphazardly taking four sub-samples (1 cm surface sediment cores with a diameter of approximately 2 cm). One sample from the sterilized commercial sediment was obtained as control for ELLS contamination with environmental DNA (Torti et al., 2015) and sample collection (Hornung et al., 2019).

Bacterioplankton communities were sampled by filtering 250 ml of water through a 0.22-μm pore size polycarbonate membrane using a EZ-Fit™ vacuum filtration system (Millipore). One specimen of each reef organism was sampled from each microcosm. After sampling, the organisms were rinsed with sterilized artificial seawater (filtered through 0.22-μm pores) and removed from their carbonate stones. All sediment samples, whole membrane filters and reef organisms were frozen at -80 °C until DNA extraction. An overview of the sample metadata can be found in supplementary table 1.

PCR-ready genomic DNA was isolated from all samples using the FastDNA® SPIN soil Kit (MP biomedicals) following the manufacturer’s instructions. Briefly, the whole membrane filter (bacterioplankton communities) and sediment (∼500 mg) and host organisms were transferred to Lysing Matrix E tubes containing a mixture of ceramic and silica particles. Due to the small size of *Zoanthus* sp. (< 60 mg) and *Chondrilla* sp. (varied between 130 and 500 mg), whole organisms were used for extraction. For *M. digitata* and *M. capricornis*, fragments (tissue and skeleton) (∼500 mg) were first snap frozen in liquid nitrogen and subsequently ground in a heat-sterilized mortar and pestle. Blank negative controls (without samples) were also included to evaluate sample contamination during DNA extraction. Microbial cell lysis was performed in the FastPrep Instrument (MP biomedicals) for 2 × 40 seconds at a speed setting of 6.0 ms^-1^. Extracted DNA was eluted in 50 μl of DNase/pyrogen-free water and stored at -20 °C until further use.

#### 2.3.2. 16S rRNA gene library preparation and sequencing

The V3/V4 variable region of the 16S rRNA gene was amplified using primers 341F 5’CCTACGGGNGGCWGCAG′3 and 785R 5′GACTACHVGGGTATCTAATCC′3 (Klindworth et al., 2013) with Illumina Nextera XT overhang adapters for a dual-PCR library preparation approach. PCRs were performed using 1 to 3 μl of DNA template, 10 μl of HS ReadyMix (KAPA HiFi Roche), and 0.6 μl of the forward and reverse primers in a concentration of 10 pmol/μl. Reaction mixes were finalized by the addition of mQ water (Ultrapure) to a final volume of 20 μl. The PCR conditions consisted of initial denaturing at 95 °C for 3 min, followed by 30 cycles of 98 °C for 20 s, 57 °C for 30 s, and 72 °C for 30 s, after which a final elongation step at 72 °C for 1 min was performed. We checked for the success of amplification, relative intensity of the bands, and contamination using 2% Invitrogen E-gels with 3 μl of PCR product.

PCR products were cleaned with magnetic beads at a ratio of 0.9:1 using a magnetic extractor stamp, after which a second PCR was performed. The 25 μl reaction mix consisted of 4 μl of the first PCR product, 12.5 μl HS ReadyMix (KAPA HiFi Roche), 2 × 1 μl (concentration of 10 pmol/μl) MiSeq Nextera XT adapters (dual indexed, Illumina), and 6.5 μl of mQ water (Ultrapure). PCR conditions consisted of initial denaturing at 95 °C for 3 min, followed by 8 cycles of 98 °C for 20 s, 55 °C for 30 s, and 72 °C for 30 s, after which a final elongation step at 72 °C for 5 min was performed. DNA molarity and fragment sizes of the PCR products were measured on a fragment analyzer 5300 (Agilent) and subsequently, each 96-well plate was normalized and pooled together in subpools using the Qiagen QIAgility. The DNA molarity of the subpools was measured on the Agilent 4150 TapeStation to combine the subpools into a final pool. The pool of normalized DNA was cleaned one last time using magnetic beads at a ratio of 0.65:1 and thereafter sequenced at a commercial company (Baseclear, Leiden, The Netherlands) on the Illumina MiSeq platform using 2 × 300 bp paired-end sequencing (Illumina MiSeq PE300). Three negative control samples were included to detect possible contamination during library preparation and sequencing. Sequences from each end were paired following Q25 quality trimming and removal of short reads (<150 bp). The DNA sequences generated in this study can be downloaded from NCBI BioProject Ids: PRJNA904682.

#### 2.3.3. Bacterial Community Analysis

Demultiplexed FASTQ files, which contained paired forward and reverse reads for each sample, were imported and visualized using QIIME2 (Bolyen et al., 2019). Subsequently, forward and reverse sequences were trimmed to a length of 245 and 200 nt, respectively, using the DADA2 plugin (Callahan et al., 2016). The DADA2 analysis produced a quality filtered table of all operational taxonomic units (OTUs), a fasta file of representative sequences, and a table summarising the denoising statistics. Using the described method, the OTUs generated are equal to exact amplicon sequence variants (ASVs). Following this, the QIIME2 feature-classifier plugin with the extract-reads option was used to extract reads from the Silva database with the silva-138-99-seqs.qza file as input and the forward and reverse PCR primers as parameters. This produced a file of reference sequence reads, which was used as input for the feature-classifier plugin with the fit-classifier-naive-bayes option. The feature-classifier plugin was then used with the classify-sklearn method and the i-reads argument set to the representative sequences file generated by the DADA2 analysis, which produced a table with taxonomic assignment for all OTUs. Mitochondria, chloroplasts, and Eukaryota were filtered out from the obtained OTU table using the QIIME2 taxa plugin with the filter-table method. The OTU count table is presented in supplementary table 2.

OTUs classified as Archaea, unassigned at the phylum level and that occurred in the triple-autoclaved commercial sediment (control for sampling and eDNA contamination) and negative controls were removed. All OTUs removed following detection in the control samples are listed in supplementary table 3. Overall, the removed OTUs were assigned to known contaminants, for example, the genera *Ralstonia, Burkholderia-Caballeronia-Paraburkholderia, Reyranella, Bacillus* and *Bradyrhizobium* (Weyrich et al., 2019). After these filtering steps, we removed nine samples with low (< 4000 reads) read counts. Among these were two sediment samples, one water sample, one sample of *M. capricornis* and six samples of *Zoanthus sp*. The low sequence quality obtained from *Zoanthus* sp. resulted in a significant loss of replicates, and therefore, we decided to exclude this organism from further bacterial community analysis.

The ten most abundant OTUs of each biotope were referenced against the NCBI nucleotide database using NCBI Basic Local Alignment Search Tool (BLAST) (Boratyn et al., 2013). BLAST identifies locally similar regions between sequences, compares sequences to extant databases and assesses the significance of matches; functional and evolutionary relationships can subsequently be inferred.

The DNA sequences generated in this study can be downloaded from NCBI BioProject Ids: PRJNA904682.

### 2.4. Statistical analyses

We checked for deviations from normality in Fv/Fm ratios with the shapiro.test() function and tested for homogeneity of variance with the bartlett.test() function in R (http://www.r-project.org/; Accessed June 2022). We tested for significant differences among treatments in normally distributed Fv/Fm ratios (*M. capricornis, S. glaucum*) with a one-way ANOVA using the aov() (Stats package) function and tested for significant differences in non-normally distributed Fv/Fm ratios (*M. digitata*) with a permutational analysis of variance (PERMANOVA) using the adonis2() function (999 permutations, Vegan package) (Vegan package) in R. In the aov() function, the Fv/Fm ratio was the response variable and treatment was the independent variable. In the adonis2() analysis, the Euclidean distance matrix of Fv/Fm ratio was the response variable with treatment as independent variable. The number of permutations was set at 999.

A table containing the OTU counts was imported into R. Differences in higher taxon abundances were assessed using the glm() function in R with the family argument to “quasibinomial”.

Variation in bacterial composition among treatments was visualised with Principal Coordinates Analysis (PCO). For the PCO analysis, the OTU table was rarefied per biotope to the minimum sample size using the rrarefy() function of the R package vegan. For compositional analyses, the OTU table was log (x + 1) transformed (in order to normalise the distribution of data) and a distance matrix constructed using the Bray–Curtis index with the vegdist() function in the vegan package in R. We tested for significant differences in OTU composition among treatments within each biotope with a permutational analysis of variance (PERMANOVA) using the adonis2() function (999 permutations).

Finally, we used the boruta function in R (Kursa et al., 2010) to identify specific OTUs, which were positively or negatively associated with HS supplementation. Boruta, named after a slavic forest demon, is a random forest wrapper, which is used to evaluate feature importance.

## 3. Results

### 3.1. Photobiology of the host-organisms

HS supplementation proved to be a significant independent predictor of variation in the Fv/Fm ratios of *M. digitata, M. capricornis* and *S. glaucum* (Fig. 2, (PERM)ANOVA, *p* = 0.007, *p* < 0.001 and *p* = 0.005, respectively, supplementary table 4). In all three biotopes, the Fv/Fm ratios were higher on average under HS supplementation. Temperature was, furthermore, a significant independent predictor of the photosynthetic efficiencies of *M. digitata* and *M. capricornis* ((PERM)ANOVA, *p* = 0.031 and *p* < 0.001, respectively, supplementary table 4). In both biotopes the Fv/Fm ratios were lower on average in heat-treated microcosms. In addition to this, there was a significant interactive effect of temperature and HS supplementation on the photosynthetic efficiency of *M. digitata* (PERMANOVA, *p* = 0.008, supplementary table 4). Fv/Fm ratios were lower in heat-treated microcosms without HS supplementation, whereas there was no such effect with HS supplementation.

**Fig. 2.**
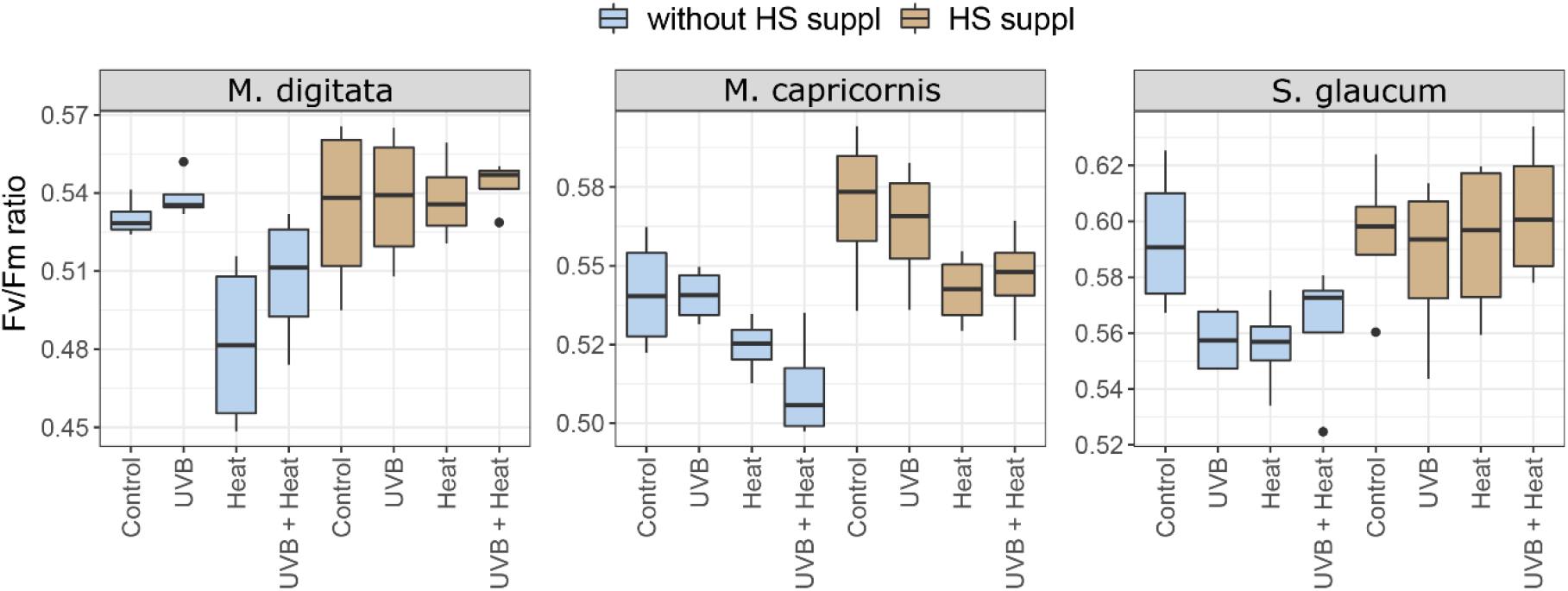
Boxplots of the maximum quantum yield of photosystem II (Fv/Fm ratio) of M. digitata, M. capricornis and S. glaucum. Fv/Fm ratios are grouped per treatment within each biotope.

### 3.2. Bacterial community analysis

The present dataset consisted of 214 samples. After quality control and removal of mitochondria, chloroplasts and OTUs unassigned at Domain and Phylum level, the data set consisted of 4,339,623 sequences and 21,068 OTUs, which were assigned to 48 phyla. The number of OTUs recorded per sample varied from 51 in a *Chondrilla* sp. sample to 2372 in a sediment sample (supplementary table 5). The most abundant (in terms of sequences) phyla sampled in the present study included Proteobacteria, Planctomycetes, Bacteroidetes, Cyanobacteria, Actinobacteria, Verrucomicrobiota, Patsecibacteria, Firmicutes, Chloroflexi and Acidobacteriota. The richest (in terms of OTUs) phyla included Proteobacteria, Planctomycetota, Bacteroidota, Verrucomicrobiota, Patescibacteria, Actinobacteria, NB1-j, Acidobacteriota, Myxococcota and Chloroflexi.

#### 3.2.1 Community composition

HS supplementation proved to be a significant predictor of variation in OTU composition of all biotopes with the exception of *M. capricornis* (PERMANOVA, *p* < 0.05, supplementary table 6). Temperature also proved a significant predictor of variation in OTU composition in water, *S. glaucum* and, *Chondrilla* sp. (PERMANOVA, *p* = 0.043, *p* = 0.028 and *p* = 0.026, respectively, supplementary table 6). Finally, there was a significant interactive effect of Heat, UVB radiation and HS supplementation in *M. digitata* (PERMANOVA, *p* = 0.039). PCO ordinations presented in Fig. 3 show that samples subjected to HS supplementation clearly separated along the first two axes. For *M. digitata*, the ordination, furthermore, showed a separate cluster of Heat and UVB-treated samples in microcosms not supplemented with HS.

**Fig. 3.**
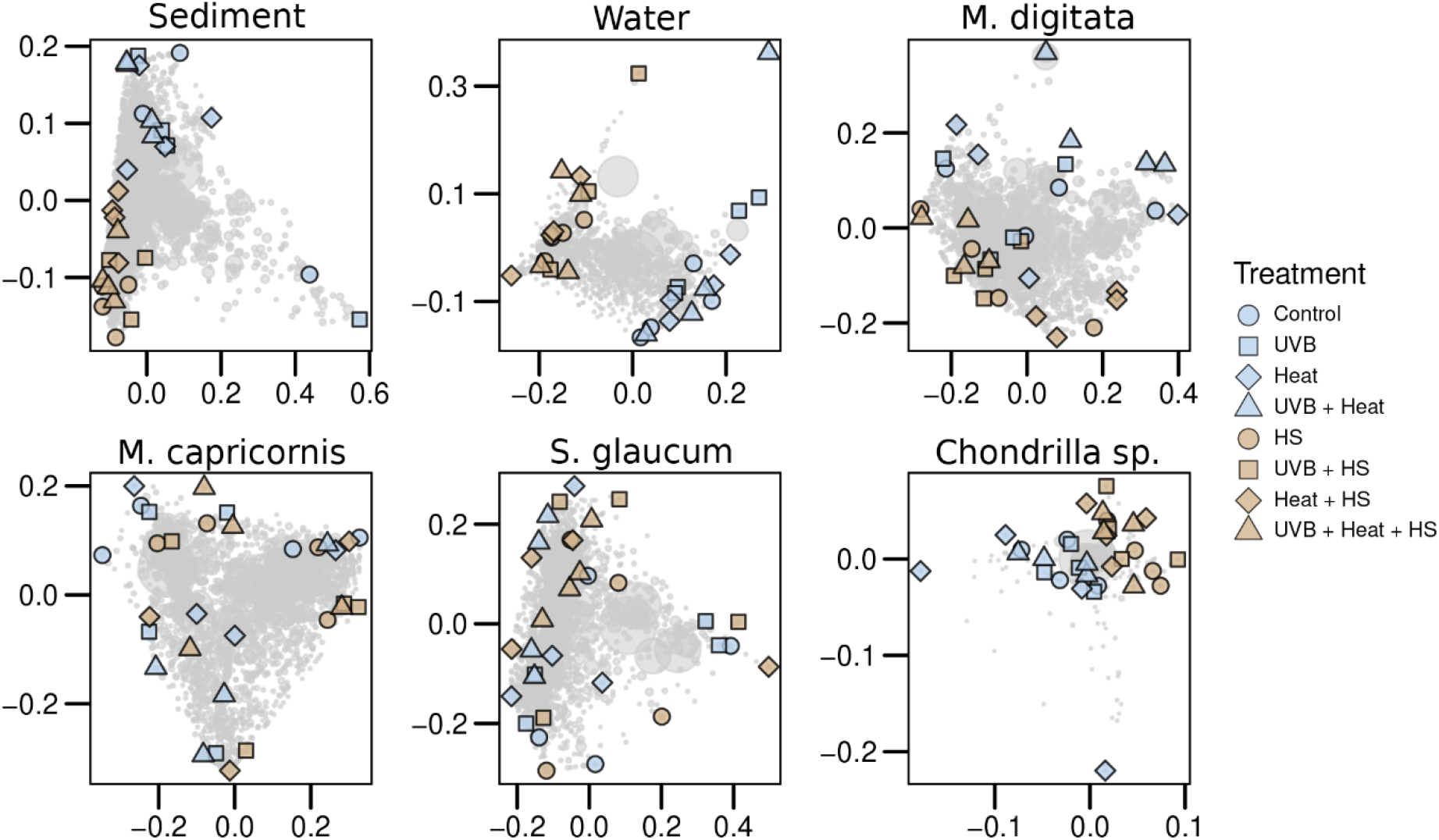
Ordinations showing the first two axes of the principal coordinates analysis (PCO) of prokaryotic OTU composition of the studied biotopes. The PCO was generated using the cmdscale() function in the R base package and wascores() function in vegan. Prior to the PCO, the raw data were log(x + 1) transformed and used to produce a distance matrix based on the Bray–Curtis distance with the vegdist() function in vegan (Oksanen et al., 2012). Light grey symbols represent operational taxonomic unit (OTU) scores with the symbol size proportional to their abundance (number of sequence reads). The percentage of variation explained by the first two axes was 21.37% in sediment, 27.64% in water, 18.38% in M. digitata, 28.16% in M. capricornis, 24.7% in S. glaucum and 32.84% in Chondrilla sp..

#### 3.2.2. Higher taxon abundance

The classes Gammaproteobacteria and Bacteroidia had significantly higher, and the orders Rhodobacterales, Pirellulales and Microtrichales significantly lower relative abundances in HS-supplemented microcosms (GLMs, *p* < 0.05, supplementary Fig. 1, Fig. 4 and supplementary table 6). In *Chondrilla* sp., the relative abundances of the BD2-11 terrestrial group and HOC36 were significantly higher and lower, respectively. In *M. digitata*, the relative abundance of the order Pirellulales was significantly higher with HS supplementation. The relative abundance of the order Cytophagales was significantly lower in heat treated microcosms with UVB and without HS than in HS-supplemented microcosms without UVB. In *S. glaucum*, there was a significant independent effect of temperature on the relative abundances of the classes Firmicutes and Bacilli and order Entoplasmatales, all lower with heat, and the order Rhodobacterales, higher with heat. In water, the ABY1 order was significantly more abundant in HS-supplemented microcosms, the order Rhodobacterales in heat-treated microcosms and the order Caulobacterales in HS-supplemented and heat-treated microcosms, but this effect was not apparent under UVB exposure. No significant differences were observed for the four most abundant classes and orders in *M. capricornis*.

**Fig. 4.**
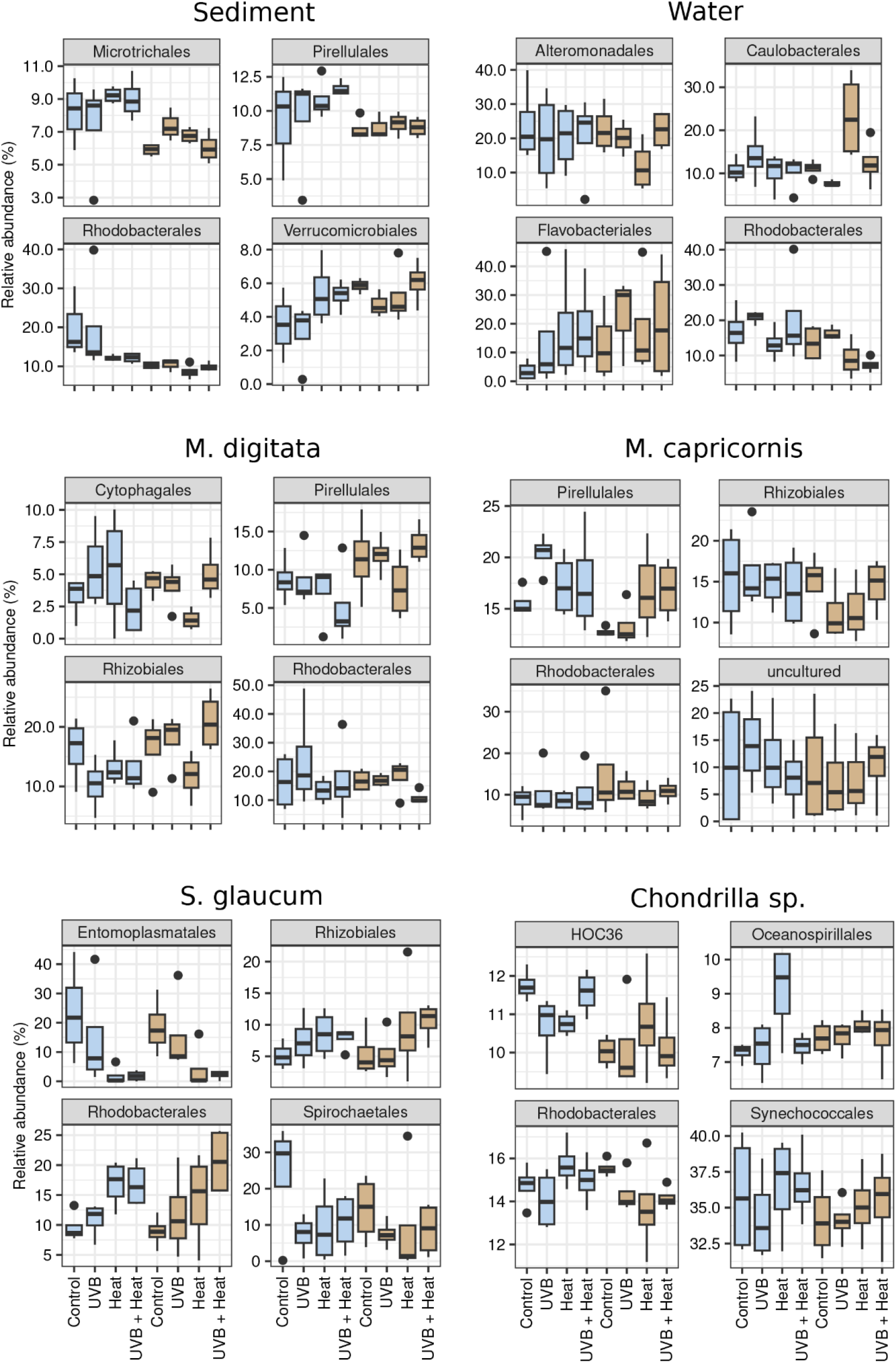
Boxplots of the relative abundance of the four most abundant orders in sediment, water, M. digitata, M. capricornis, S. glaucum and Chondrilla sp. under the independent and combined effects of UVB, Heat and HS supplementation. Relative abundances are grouped per treatment within each biotope. Boxplots of treatments without HS supplementation are depicted in blue, boxplots of treatments with HS supplementation are depicted in brown.

#### 3.2.3. Abundant OTUs

The ten most abundant OTUs in sediment, water, *M. digitata, S. glaucum*. and *Chondrilla* sp. are shown in Fig. 5 with and OTUs highlighted, which contributed to the significant compositional differences described above. In line with the significant impact of HS on sediment composition, we observed that OTU-11, assigned to family Rhodobacteraceae and closely related to an organism detected in the coral *Porites cylindrica* (100% sequence similarity), was particularly abundant in microcosms without HS supplementation (supplementary table 7). In contrast, OTUs 17 and 66, assigned to the genera *Filomicrobium* and *Halieaceae*, respectively, were particularly abundant in sediment of HS-supplemented microcosms.

**Fig. 5.**
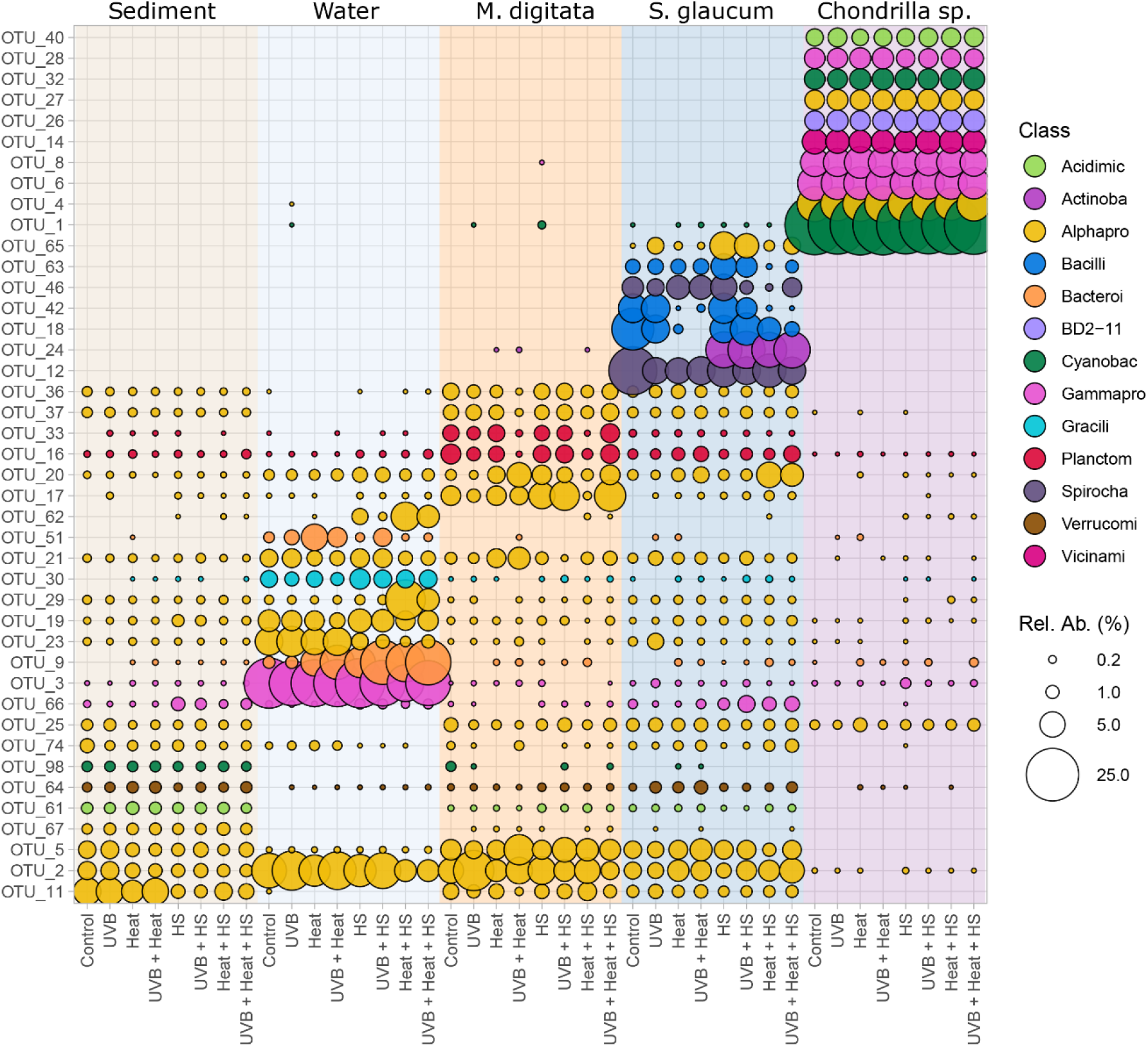
Mean relative abundances of the ten most abundant OTUs in sediment, water, M. digitata, S. glaucum and Chondrilla sp.. Relative abundances are grouped per treatment within each biotope. Symbols are proportional to the relative abundance of the respective OTU and colour-coded following their class-level taxonomic assignment.

In line with the significant impact of HS on water bacterial composition, we found that OTU-23, assigned to the genus *Maricaulis* (family Hyphomonadaceae), was particularly abundant in microcosms without HS supplementation. OTU-62, in turn, also assigned to the Hyphomonadaceae family, was particularly abundant in HS-supplemented microcosms. Additionally, OTU-9, assigned to the NS3a marine group, was markedly more abundant in heat-treated and HS-supplemented microcosms. OTU-29, assigned to the family Hyphomonadaceae, was particularly abundant in heat treated microcosms with HS supplementation. These previously mentioned OTUs were all closely related to organisms previously detected or isolated from seawater samples (100% sequence similarities, supplementary table 7).

OTU-66, assigned to the genus *Halieaceae*, occurred in all treatments with HS supplementation, but was absent from heat, and heat + UVB treated microcosms without HS supplementation. This is in line with the significant interaction of HS supplementation, heat and UVB on the bacterial composition of *M. digitata*. OTUs 36 and 37, assigned to the genera *Methyloceanibacter* and *Filomicrobium*, in turn, had markedly lower relative abundances in heat + UVB treated microcosms, but HS supplementation appeared to mitigate this effect.

In consonance with the significant effects of temperature and HS supplementation on the bacterial composition of *S. glaucum*, OTU 24, assigned to the genus *Rhodococcus*, was highly abundant in microcosms with and absent from microcosms without HS supplementation. Interestingly, this OTU was closely related to a humus utilizing *Rhodococcus sp. strain* isolated from marine sediment (100% sequence similarity; GenBank MT626344.1). OTUs 18 and 42, both assigned to the genus candidatus *Hepatoplasma*, were highly abundant under ambient temperatures, but markedly less so in heat-treated microcosms. In contrast, OTUs 2 and 20, assigned to the alphaproteobacterial families Rhodobacteraceae and Methyloligellaceae, were markedly more abundant in heat-treated microcosms. OTU-2 was closely related to a *Tritonibacter litoralis* isolate, and OTU 20 to an organism obtained from the sponge *Terpios Hoshinota* (both 100% sequence similarity; GenBank MN544912.1 and KX177464.1, respectively).

In contrast to the previously described biotopes, there appeared to be a much higher degree of stability in the bacterial community of *Chondrilla* sp., suggesting that the significant impact of HS on bacterial composition affected the less abundant OTUs. Most of these OTUs were, furthermore, highly similar to organisms previously obtained from other sponges, e.g., *Haliclona sp* and *Terpios hoshinota* (supplementary table 7).

#### 3.2.4. Significant predictor OTUs

Significant predictor OTUs based on the Boruta analysis and which had an importance level > 5.00% are presented in Fig. 6 for sediment, water, *M. digitata, S. glaucum* and *Chondrilla* sp.. A number of OTUs exhibited consistent responses to HS supplementation across biotopes (sediment, water, *M. digitata* and *Glaucum* sp.). OTUs 39, 43, 81, and 310, for example, were all more abundant in microcosms without HS supplementation, whereas OTUs 50, 66, and 256 were all more abundant in microcosms with HS supplementation (see supplementary table 8 for statistical results).

**Fig. 6.**
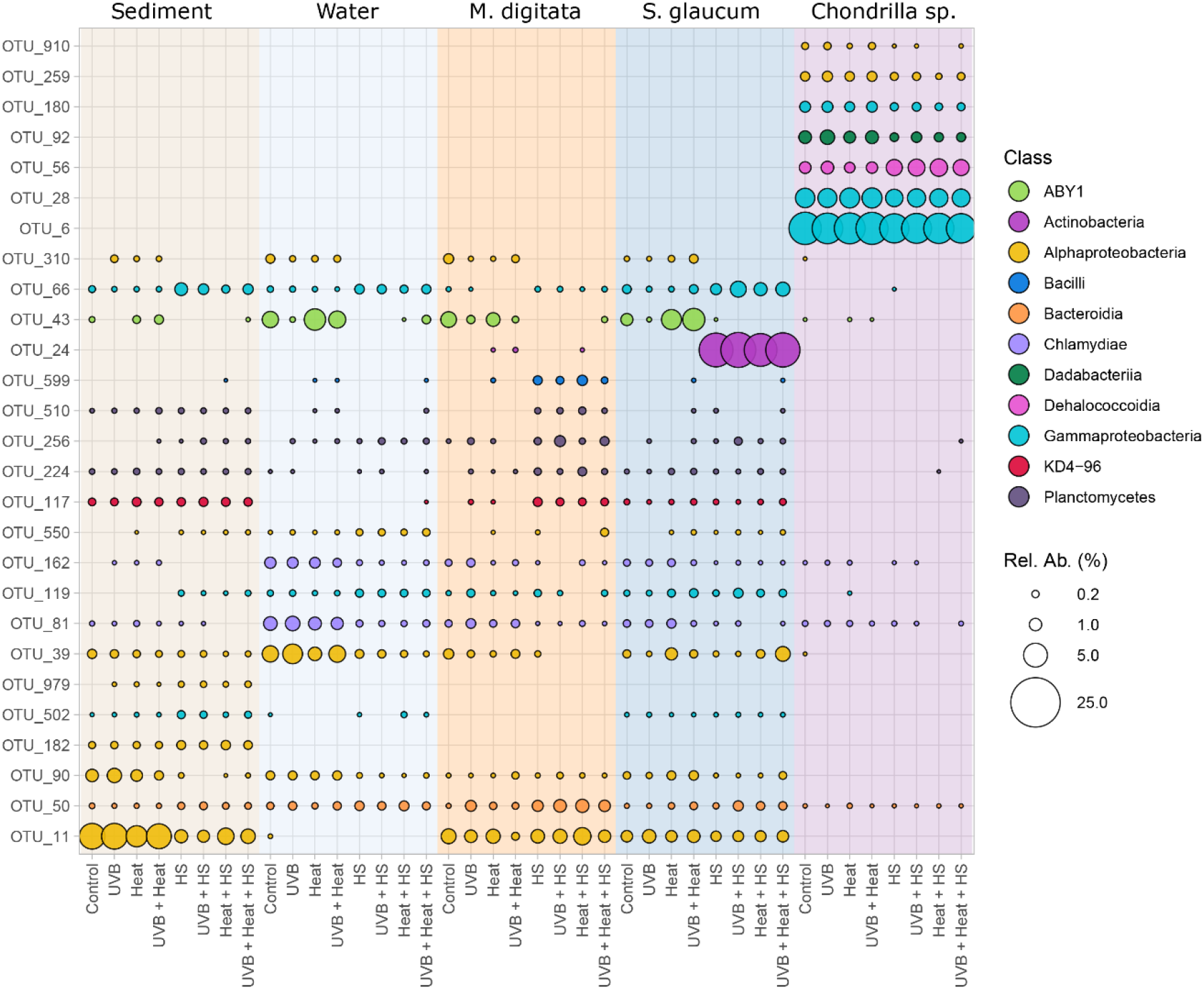
Mean relative abundances of the OTUs with more than 5.00% importance in the Boruta analysis in sediment, water, M. digitata, S. glaucum and Chondrilla sp.. Relative abundances are grouped per treatment within each biotope. Symbols are proportional to the relative abundance of the respective OTU and colour-coded following their class-level taxonomic assignment.

## 4. Discussion

### 4.1. Effects of humic substances, temperature and UVB on photosynthetic activity

The adverse effects of elevated temperature and UVB radiation on coral health have been well documented (Hughes et al., 2017). Various studies, however, have observed considerable site-specific variation to these stressors (Graham et al., 2011; Hughes et al., 2003; Hughes et al., 2010). One explanation for this observed heterogeneity lies in the role that local environmental conditions play in influencing host coral resilience (Montefalcone et al., 2020). Poor water quality and increased fishing activities, for example, have been suggested to exacerbate climate-induced reef decline (MacNeil et al., 2019). However, much less is known concerning possible mitigating factors (Wyatt et al., 2020). In the current study, we proposed that terrestrially-derived humic substances might play a mitigating role in the responses of coral reef organisms and associated bacterial communities to stress-inducing temperatures and UVB radiation.

The photo-system type II (PSII) maximum potential quantum efficiency of the coral-associated zooxanthellae is widely used as a quantitative indicator of coral photo-oxidative stress (Downs et al., 2002; Travesso et al., 2023). According to the “oxidative stress theory”, temperature and UVB-induced coral bleaching result from a PSII-dysfunction in the zooxanthellae chloroplasts leading to reduced photosynthetic efficiency, excessive production of reactive oxygen species (ROS) and deactivation of various ROS-neutralizing pathways. ROS can subsequently accumulate and diffuse into the coral-host tissue causing oxidative stress and result in zooxanthellae expulsion (Downs et al., 2002). In the present study, we found that elevated temperatures were associated with significantly lower Fv/Fm ratios in the hard corals *M. digitata* and *M. capricornis*, but not in the soft coral *S. glaucum*. The results for both hard coral species confirm previous laboratory studies and in situ observations (Higuchi et al., 2013; Manullang et al., 2020) and suggest that the photosynthetic machinery of the coral symbionts were damaged by the high-temperature treatment (Downs et al., 2002). In contrast to our results, another study found a significant reduction in Fv/Fm ratios in *S. glaucum*, following exposure to a laboratory-induced heatwave (Travesso et al., 2023). Travesso et al., (2023), however, exposed their specimens for a period of ten days to an elevated temperature in contrast to the five days of our study. The lack of a significant response of *S. glaucum* in our study might, therefore, be related to the shorter duration of our experiment. Although non-significant, Fv/Fm ratios of *S. glaucum* in heat-treated microcosms without HS supplementation were lower than the controls and longer exposure might have resulted in additional reductions in photosynthetic efficiency.

In contrast to heat, HS supplementation alone had a significant and beneficial effect on Fv/Fm ratios, and, most importantly, HS supplementation appeared to mitigate the adverse effect of elevated temperature in all three tested species. Independent effects of HS supplementation on coral photosynthetic efficiency can be explained by their optical properties, which are characterized by a particularly high absorbance of wavelengths in the UV spectrum (Del Vecchio & Blough, 2004). In HS-supplemented microcosms, the corals might have increased their photosynthetic efficiency (a process called photo-acclimation) as a response to the altered light regime (Roth, 2014). Previous studies have demonstrated that elevated temperatures and intense UVB levels interact to adversely affect coral health and induce bleaching (Ferrier-Pagès et al., 2007; Fournie et al., 2012). Our results suggest that HS are able to mitigate the adverse effects of both stressors. Our findings are in accordance with previous observations of higher coral resistance to temperature-induced bleaching in reefs exposed to high CDOM concentrations (MacNeil et al., 2019). The major source of CDOM in coastal environments are soil leachates, of which the predominant components are HS (Coble, 2013).

### Effects of humic substances, temperature and UVB on environmental and host-associated bacterial communities

As previously discussed (Stuij et al., 2023), the bacterial communities present in the control microcosms were similar to natural-type communities of coral reef sediment, water and host organisms. In the present study, HS supplementation and heat were significantly associated with compositional differences in host-associated and environmental bacterial communities. Although there was no significant independent effect of UVB radiation on bacterial composition, there was a significant interaction between HS supplementation, heat and UVB radiation in the coral *M. digitata*. In the PCO analysis, *M. digitata* samples in microcosms under the combined effect of heat and UVB were distant from the controls, whereas this effect was diminished with HS supplementation.

Heat had a significant and independent effect on the bacterial community composition of the soft coral *S. glaucum* and sponge *Chondrilla* sp.. Previously, a similar response to elevated temperature was observed in the sponge *Xestospongia muta* and the coral *Acropora tenuis* (Littman et al., 2010; Lesser et al., 2016), but not in the sponge *Rhopaloeides odorabile* or the soft coral *Lobophytum pauciflorum* (Fan et al., 2013; Wessels et al., 2017).

HS supplementation significantly affected the bacterial composition of all studied biotopes, with the exception of *M. capricornis*. This is in line with previous studies that observed an effect of HS on marine bacterioplankton and fish-associated bacterial communities (Lindh et al., 2015; Louvado et al., 2021), and supports a strong modulatory effect across a broad range of marine biotopes. Several OTUs were positively or negatively associated with HS supplementation across multiple biotopes. OTUs associated with HS supplementation were assigned to a range of taxa including the gammaproteobacterial family *Halieaceae*, Interestingly, Shore et al. (2021) previously observed an enrichment of *Halieaceae* members in bacterial communities of the sponges *Xestospongia muta* and *Agelas clathrodes* following increased terrestrial run-off. In *M. digitata*, OTUs assigned to the families *Gimesiaceae* and *Pirellulaceae (*Planctomycetales), were also associated with HS supplemention. Members of the order Pirellulales have frequently been detected in association with coral and sponge hosts (Mohamed et al., 2010; Sun et al., 2022). Moreover, in a study which evaluated the bacterial communities of 26 coral genera, OTUs assigned to the order Pirellulales were mostly associated with healthy coral microbiomes, and their abundances were lower in bleached individuals (Sun et al., 2022).

In *Chondrilla* sp., an OTU, assigned to the SAR202 clade, was more abundant in HS-supplemented microcosms. SAR202 clade members have the genomic potential to synthesize a variety of secondary metabolites and co-factors, as well as to degrade recalcitrant compounds (Bayer et al., 2018). Their genomes, furthermore, included features relevant to symbiosis, such as gene clusters encoding for CRISPR-Cas systems and eukaryote-like repeat proteins (Bayer et al., 2018). In *S. glaucum*, an abundant OTU assigned to the genus *Rhodococcus*, was a significant predictor of HS supplementation, and was completely absent in microcosms without HS supplementation. *Rhodococcus* members are widely distributed in the environment and have also been observed in association with coral hosts (Hackbusch et al., 2020; Ramaprasad et al., 2018); they are known for their ability to degrade a wide range of complex organic compounds (Hackbusch et al., 2020). Several *Rhodococcus* strains obtained from tundra soils were identified as putative HS-degraders (Park et al., 2021). OTU-24, furthermore, had 100% similarity with a strain classified as *Rhodococcus* and described as a “humus-utilizing bacteria” isolated from sediment in the Pacific Ocean (Acc. nr.: MT626344.1). Various *Rhodococcus* strains from the coral species *Coscinaraea columna, Platygyra daedalea* and *Porites harrisoni* exhibited antibiotic activity against the putative pathogenic bacteria *Staphylococcus aureus, Bacillus subtilis* and *Escherichia coli* (Mahmoud & Kalendar, 2016), indicating a potential role in host defense.

In *M. digitata*, we observed a significant interactive effect of elevated temperature, UVB radiation and HS supplementation on bacterial community composition and higher taxon abundance. Several abundant OTUs were less abundant in heat and UVB-treated microcosms without HS, but this effect was not apparent under HS supplementation. This included OTUs assigned to the genera *Methyloceanibacter* and *Filomicrobium*, which have been previously identified as core members of several coral species of the genus *Montipora* (Cai et al., 2018). In contrast, an OTU assigned to the genus *Pyruvatibacter* was more abundant in heat and UVB treated microcosms without HS supplementation. Recently, genomic analysis of a *Pyruvatibacter* strain isolated from a marine microalga revealed that it could utilize various antioxidants to deal with oxidative stress (Rong et al., 2021). Together with the results of our Fv/Fm analysis, these findings strongly suggest that *M. digitata* experienced oxidative stress in the microcosms with elevated temperatures and UVB radiation and this was associated with shifts in its bacterial community; HS supplementation, however, appeared to mitigate this.

## 5. Conclusion and implications for coral reef health

In summary, our results showed that elevated temperature and UVB affected the photosynthetic activities of all corals. Importantly, the adverse effects of these abiotic factors were mitigated by HS supplementation. With respect to bacterial composition, our results showed a pronounced effect of HS across all biotopes with the possible exception of *M. capricornis*. Temperature, in contrast, only had a significant independent effect on *S. glaucum* and the sponge *Chondrilla* sp. and a significant interactive effect on *M. digitata*. Our findings indicate that HS significantly modulates coral reef bacterial communities, and may contribute to improved reef resilience to climate-related perturbations.

Periodic bleaching during severe ENSO-induced warming events has induced large-scale destruction of coral reef habitat (Hughes et al., 2017). Coral reef resilience has been suggested to play a key role in bleaching susceptibility (Graham et al., 2011; Hughes et al., 2010). Importantly, resilience is largely determined by site-specific conditions, for example, inputs of nutrients, pollutants, turbidity, sedimentation rates, and grazing activity (Babcock & Davies, 1991; MacNeil et al., 2019; Mumby et al., 2006; Zweifler et al., 2021). An improved understanding of the interplay between these different environmental factors is crucial to successfully restore and protect coral reefs. The reported effects of HS suggest that natural forests and wetlands (the predominant sources of HS inputs into coastal ecosystems) may have an important, but understudied, role in promoting coral reef resilience. As such, large-scale clearance of coastal and riparian forests may have significantly contributed to the increased vulnerability of certain coral reefs to climate-induced bleaching. Management efforts, with a focus on restoring natural forests on previously cleared lands will not only restore habitat for terrestrial organisms, but may also lead to a cascade of beneficial effects for adjacent marine ecosystems.

## Authorship contribution statement

**T.M Stuij**: Investigation, data analysis, writing –original draft, review & editing; **D.F.R. Cleary**: Supervision, conceptualization, methodology, data analysis, writing – original draft, review & editing; **N.J. de Voogd**: Supervision, conceptualization, resources, review & editing; **R.J.M. Rocha**: Resources, conceptualization; **A.R.M. Polonia**: Investigation, review & editing; **D.A.M. Silva**: Conceptualization, investigation, review & editing; **J.C. Frommlet**: Resources, review & editing; **A. Louvado**: Conceptualization, methodology, investigation, review & editing; **Y. M. Huang**: Resources; **N. van der Windt**: Investigation; **N.C.M. Gomes**: Supervision, conceptualization, resources, investigation, methodology, writing – original draft, review & editing.

## Funding

This study is funded by 4D-REEF (www.4d-reef.eu). 4D-REEF receives funding from the European Union’s Horizon 2020 research and innovation program under the Marie Sklodowska-Curie grant agreement No 813360. Additional funding was received from the Ministry of Science and Technology, Taiwan to Y. M. Huang (MOST 110-2621-B-346-001). Ana R.M. Polónia was supported by a postdoctoral scholarship (SFRH/BPD/117563/2016) funded by the Portuguese Foundation for Science and Technology (FCT)/ national funds (MCTES) and by the European Social Fund (ESF)/EU. Jörg C. Frommlet was supported by national funds (OE), through FCT, in the scope of the framework contract foreseen in the numbers 4, 5 and 6 of the article 23, of the Decree-Law 57/2016, of August 29, changed by Law 57/2017, of July 19. We acknowledge financial support to CESAM by FCT/MCTES (UIDP/50017/2020+UIDB/50017/2020+ LA/P/0094/2020), through national funds. Davide A.M. Silva was supported by a PhD grant (2020.05774.BD) funded by the Portuguese Foundation for Science and Technology (FCT), national funds (MCTES), and the European Social Fund POPH-QREN programme. Additional funding was received from the research programme NWO-VIDI with project number 16.161.301, which is financed by the Netherlands Organisation for Scientific Research (NWO).

## Declaration of competing interest

The authors declare that they have no known competing financial interests or personal relationships that could have appeared to influence the work reported in this paper.

## Data availability

Sequences generated in this study can be downloaded from the NCBI Sequence Read Archive under the BioProject accession number PRJNA904682.

## Acknowledgements

We are grateful to F. Coelho and V. Oliveira for their help during lab work and FOLEX COATING GMBH in Germany for providing samples of their specialized polyester film. The research was conducted at the Laboratory of Molecular Studies and Marine Environment, situated at the Centre for Environmental and Marine Studies (CESAM), University of Aveiro, Portugal.

## Notes

https://www.ncbi.nlm.nih.gov/bioproject/?term=PRJNA904682

### Competing Interest Statement

The authors have declared no competing interest.

## References

Ayoub, L. M. L. M., Hallock, P., Coble, P. G. P. G., & Bell, S. S. (2012). MAA-like absorbing substances in Florida Keys phytoplankton vary with distance from shore and CDOM: Implications for coral reefs. Journal of Experimental Marine Biology and Ecology, 420–421, 91–98. https://doi.org/10.1016/j.jembe.2012.03.026

Babcock, R., & Davies, P. (1991). Effects of sedimentation on settlement of Acropora millepora. Coral Reefs, 9(4), 205–208. https://doi.org/10.1007/BF00290423

Bayer, K., Jahn, M., Slaby, B., Moitinho-Silva, L., & Hentschel, U. (2018). Marine sponges as Chloroflexi hot-spots: Genomic insights and high resolution visualization of an abundant and diverse symbiotic clade. MSystems, 3, e00150–18. https://doi.org/10.1101/328013

Blackwood, J. C., Hastings, A., & Mumby, P. J. (2012). The effect of fishing on hysteresis in Caribbean coral reefs. Theoretical Ecology, 5(1), 105–114. https://doi.org/10.1007/s12080-010-0102-0

Bolyen, E., Rideout, J. R., Dillon, M. R., Bokulich, N. A., Abnet, C. C., Al-Ghalith, G. A., Alexander, H., Alm, E. J., Arumugam, M., Asnicar, F., Bai, Y., Bisanz, J. E., Bittinger, K., Brejnrod, A., Brislawn, C. J., Brown, C. T., Callahan, B. J., Caraballo-Rodríguez, A. M., Chase, J., … Caporaso, J. G. (2019). Reproducible, interactive, scalable and extensible microbiome data science using QIIME 2. Nature Biotechnology, 37(8), 852–857. https://doi.org/10.1038/s41587-019-0209-9

Boratyn, G. M., Camacho, C., Cooper, P. S., Coulouris, G., Fong, A., Ma, N., Madden, T. L., Matten, W. T., McGinnis, S. D., Merezhuk, Y., Raytselis, Y., Sayers, E. W., Tao, T., Ye, J., & Zaretskaya, I. (2013). BLAST: a more efficient report with usability improvements. Nucleic Acids Research, 41, 29–33. https://doi.org/10.1093/nar/gkt282

Bourne, D. G., Morrow, K. M., & Webster, N. S. (2016). Insights into the Coral Microbiome: Underpinning the Health and Resilience of Reef Ecosystems. Annual Review of Microbiology, 70, 317–340. https://doi.org/10.1146/annurev-micro-102215-095440

Burkepile, D. E., Shantz, A. A., Adam, T. C., Munsterman, K. S., Speare, K. E., Ladd, M. C., Rice, M. M., Ezzat, L., McIlroy, S., Wong, J. C. Y., Baker, D. M., Brooks, A. J., Schmitt, R. J., & Holbrook, S. J. (2020). Nitrogen Identity Drives Differential Impacts of Nutrients on Coral Bleaching and Mortality. Ecosystems. 23, 798–811 https://doi.org/10.1007/s10021-019-00433-2

Cai, L., Tian, R. M., Zhou, G., Tong, H., Wong, Y. H., Zhang, W., Chui, A. P. Y., Xie, J. Y., Qiu, J. W., Ang, P. O., Liu, S., Huang, H., & Qian, P. Y. (2018). Exploring coral microbiome assemblages in the South China Sea. Scientific Reports, 8(1), 1–13. https://doi.org/10.1038/s41598-018-20515-w

Callahan, B. J., McMurdie, P. J., Rosen, M. J., Han, A. W., Johnson, A. J. A., & Holmes, S. P. (2016). DADA2: High-resolution sample inference from Illumina amplicon data. Nature Methods, 13(7), 581–583. https://doi.org/10.1038/nmeth.3869

Cleary DFR, Polónia ARM, Renema W, Hoeksema BW, Wolstenholme J, Tuti Y, de Voogd NJ. 2014. Coral reefs next to a major conurbation: a study of temporal change (1985-2011) in coral cover and composition in the reefs of Jakarta Indonesia. Marine Ecology Progress Series. 501: 89–98. doi: 10.3354/meps10678.

Cleary DFR, Polónia ARM, Renema W, Hoeksema BW, Rachello-Dolmen PG, Moolenbeek RG, Budiyanto A, Yahmantoro Tuti Y, Giyanto Draisma SG, Prud’homme van Reine WF, Hariyanto R, Gittenberger A, Rikoh MS, de Voogd NJ. 2016. Variation in the composition of corals, fishes, sponges, echinoderms, ascidians, molluscs, foraminifera and macroalgae across a pronounced in-to-offshore environmental gradient in the Jakarta Bay-Thousand Islands coral reef complex. Marine Pollution Bulletin 110: 701–17. DOI: 10.1016/j.marpolbul.2016.04.042.

Coble, P. G. (2013). Colored dissolved organic matter in seawater. In Subsea Optics and Imaging. Woodhead Publishing Limited. https://doi.org/10.1533/9780857093523.2.98

Coelho, F. J. R. C., Rocha, R. J. M., Pires, A. C. C., Ladeiro, B., Castanheira, J. M., Costa, R., Almeida, A., Cunha, Â., Lillebø, A. I., Ribeiro, R., Pereira, R., Lopes, I., Marques, C., Moreira-Santos, M., Calado, R., Cleary, D. F. R., & Gomes, N. C. M. (2013). Development and validation of an experimental life support system for assessing the effects of global climate change and environmental contamination on estuarine and coastal marine benthic communities. Global Change Biology, 19(8), 2584–2595. https://doi.org/10.1111/gcb.12227

Crain, C. M., Halpern, B. S., Beck, M. W., & Kappel, C. V. (2009). Understanding and managing human threats to the coastal marine environment. Annals of the New York Academy of Sciences, 1162(1), 39–62.

da Silva, M. S. R. de A., Huertas Tavares, O. C., Ribeiro, T. G., da Silva, C. S. R. de A., da Silva, C. S. R. de A., García-Mina, J. M., Baldani, V. L. D., Calderín García, A., Berbara, R. L. L., & Jesus, E. da C. (2021). Humic acids enrich the plant microbiota with bacterial candidates for the suppression of pathogens. Applied Soil Ecology, 168, e104146. https://doi.org/10.1016/j.apsoil.2021.104146

Del Vecchio, R., & Blough, N. V. (2004). On the origin of the optical properties of humic substances. Environmental Science and Technology, 38(14), 3885–3891. https://doi.org/10.1021/es049912h

Doering, T., Wall, M., Putchim, L., Rattanawongwan, T., Schroeder, R., Hentschel, U., & Roik, A. (2021). Towards enhancing coral heat tolerance: a “microbiome transplantation” treatment using inoculations of homogenized coral tissues. Microbiome, 9(1), 102 https://doi.org/10.1186/s40168-021-01053-6

dos Santos, U. J., Duda, G. P., Marques, M. C., Valente de Medeiros, E., de Sousa Lima, J. R., Soares de Souza, E., Brossard, M., & Hammecker, C. (2019). Soil organic carbon fractions and humic substances are affected by land uses of Caatinga forest in Brazil. Arid Land Research and Management, 33(3), 255–273. https://doi.org/10.1080/15324982.2018.1555871

Downs, C. A., Fauth, J. E., Halas, J. C., Dustan, P., Bemiss, J., & Woodley, C. M. (2002). Oxidative stress and seasonal coral bleaching. Free Radical Biology and Medicine, 33(4), 533–543 https://doi.org/10.1016/S0891-5849(02)00907-3

Eaton, A. (1995). Measuring UV-absorbing organics: a standard method. Journal - American Water Works Association, 87(2), 86–90. https://doi.org/10.1002/j.1551-8833.1995.tb06320.x

Esham, E. C., Ye, W., & Moran, M. A. (2000). Identification and characterization of humic substances-degrading bacterial isolates from an estuarine environment. FEMS Microbiology Ecology, 34(2), 103–111 https://doi.org/10.1016/S0168-6496(00)00078-7

Fan, L., Liu, M., Simister, R., Webster, N. S., & Thomas, T. (2013). Marine microbial symbiosis heats up: The phylogenetic and functional response of a sponge holobiont to thermal stress. ISME Journal, 7(5), 991–1002. https://doi.org/10.1038/ismej.2012.165

Ferrier-Pagès, C., Richard, C., Forcioli, D., Allemand, D., Pichon, M., & Shick, J. M. (2007). Effects of temperature and UV radiation increases on the photosynthetic efficiency in four scleractinian coral species. Biological Bulletin, 213(1), 76–87. https://doi.org/10.2307/25066620

Ferrier-Pagès, Gattuso, Dallot & Jaubert. (2000). Effect of nutrient enrichment on growth and photosynthesis of the zooxanthellate coral Stylophora pistillata. Coral Reefs, 19, 103–113 https://doi.org/10.1007/s003380000078

Fichot, C.G., Kaiser, K., Hooker, S.B., Amon, R.M.W., Babin, M., Belanger, S., Walker, S.A., Benner, R., 2013. Pan-arctic distributions of continental runoff in the Arctic Ocean. Sci. Rep., 3, 1053.

Fournie, J. W., Vivian, D. N., Yee, S. H., Courtney, L. A., & Barron, M. G. (2012). Comparative sensitivity of six scleractinian corals to temperature and solar radiation. Diseases of Aquatic Organisms, 99(2), 85–93 https://doi.org/10.3354/dao02459

Gerea, M., Pérez, G. L., Unrein, F., Soto Cardenas, C., Morris, D., & Queimalinos, C. (2017). CDOM and the underwater light climate in two shallow North Patagonian lakes: evaluating the effects on nano and microphytoplankton community structure. Aquatic sciences, 79, 231–248.

Gilbert, J. A., Blaser, M. J., Caporaso, J. G., Jansson, J. K., Lynch, S. V., & Knight, R. (2018). Current understanding of the human microbiome. Nature Medicine, 24(4), 392–400. https://doi.org/10.1038/nm.4517

Glasl, B., Herndl, G. J., & Frade, P. R. (2016). The microbiome of coral surface mucus has a key role in mediating holobiont health and survival upon disturbance. ISME Journal, 10(9), 2280–2292. https://doi.org/10.1038/ismej.2016.9

Glynn, P. W. (1984). Widespread Coral Mortality and the 1982–83 El Niño Warming Event. Environmental Conservation, 11(2), 133–146. https://doi.org/10.1017/S0376892900013825

Grottoli, A. G., Wilkins, M. J., Johnston, M. D., Levas, S., Schoepf, V., Dalcin, M. P., Wilkins, M. J., Warner, M. E., Cai, W.-J., Hoadley, K. D., Pettay, D. T., & Melman, T. F. (2018). Coral physiology and microbiome dynamics under combined warming and ocean acidification. PloS One, 13(1), e0191156.

Hackbusch, S., Noirungsee, N., Viamonte, J., Sun, X., Bubenheim, P., Kostka, J. E., Müller, R., & Liese, A. (2020). Influence of pressure and dispersant on oil biodegradation by a newly isolated Rhodococcus strain from deep-sea sediments of the gulf of Mexico. Marine Pollution Bulletin, 150, 110683. https://doi.org/10.1016/j.marpolbul.2019.110683

Hall, E. R., Muller, E. M., Goulet, T., Bellworthy, J., Ritchie, K. B., & Fine, M. (2018). Eutrophication may compromise the resilience of the Red Sea coral Stylophora pistillata to global change. Marine Pollution Bulletin, 131, 701–711. https://doi.org/10.1016/j.marpolbul.2018.04.067

Higuchi, T., Agostini, S., Casareto, B. E., Yoshinaga, K., Suzuki, T., Nakano, Y., Fujimura, H., & Suzuki, Y. (2013). Bacterial enhancement of bleaching and physiological impacts on the coral Montipora digitata. Journal of Experimental Marine Biology and Ecology, 440, 54–60. https://doi.org/10.1016/j.jembe.2012.11.011

Hornung, B. V. H., Zwittink, R. D., & Kuijper, E. J. (2019). Issues and current standards of controls in microbiome research. FEMS Microbiology Ecology, 95(5), 1–7. https://doi.org/10.1093/femsec/fiz045

Hughes, T., Baird, A., Bellwood, D., Card, M., Connolly, S., Folke, C., Grosberg, R., Hoegh-Guldberg, O., Jackson, J., Kleypas, J., Lough, J., Marshall, P., Nyström, M., Palumbi, S., Pandolfi, J., Rosen, B., & Roughgarden, J. (2003). Climate change, human impacts, and the resilience of coral reefs. In Science, 301(5635), 929–933. https://doi.org/10.1126/science.1085046

Hughes, T. P., Graham, N. A. J., Jackson, J. B. C., Mumby, P. J., & Steneck, R. S. (2010). Rising to the challenge of sustaining coral reef resilience. In Trends in Ecology and Evolution, 25(11), 633–642. https://doi.org/10.1016/j.tree.2010.07.011

Hughes, T. P., Kerry, J. T., Álvarez-Noriega, M., Álvarez-Romero, J. G., Anderson, K. D., Baird, A. H., Babcock, R. C., Beger, M., Bellwood, D. R., Berkelmans, R., Bridge, T. C., Butler, I. R., Byrne, M., Cantin, N. E., Comeau, S., Connolly, S. R., Cumming, G. S., Dalton, S. J., DiazPulido, G., … Wilson, S. K. (2017). Global warming and recurrent mass bleaching of corals. Nature, 543(7645), 373–377. https://doi.org/10.1038/nature21707

Jones, R. J., & Yellowlees, D. (1997). Regulation and control of intracellular algae (= zooxanthellae) in hard corals. Philosophical Transactions of the Royal Society B: Biological Sciences, 352(1352), 457–468. https://doi.org/10.1098/rstb.1997.0033

Kegler, H. F., Hassenrück, C., Kegler, P., Jennerjahn, T. C., Lukman, M., Jompa, J., & Gärdes, A. (2018). Small tropical islands with dense human population: differences in water quality of nearshore waters are associated with distinct bacterial communities. PeerJ, 6, e4555.

Klindworth, A., Pruesse, E., Schweer, T., Peplies, J., Quast, C., Horn, M., & Glöckner, F. O. (2013). Evaluation of general 16S ribosomal RNA gene PCR primers for classical and next-generation sequencing-based diversity studies. Nucleic Acids Research, 41(1), 1–11. https://doi.org/10.1093/nar/gks808

Kursa, M. B., Jankowski, A., & Rudnicki, W. R. (2010). Boruta - A system for feature selection. Fundamenta Informaticae, 101(4), 271–285. https://doi.org/10.3233/FI-2010-288

Kusuma, D. W., Murdimanto, A., Aden, L. Y., Sukresno, B., Jatisworo, D., & Hanintyo, R. (2017, December). Sea surface temperature dynamics in Indonesia. In IOP Conference Series: Earth and Environmental Science (Vol. 98, No. 1, p. 012038). IOP Publishing.

Lesser, M. P., Fiore, C., Slattery, M., & Zaneveld, J. (2016). Climate change stressors destabilize the microbiome of the Caribbean barrel sponge, Xestospongia muta. Journal of Experimental Marine Biology and Ecology, 475, 11–18. https://doi.org/10.1016/j.jembe.2015.11.004

Li, Y., Fang, F., Wei, J., Wu, X., Cui, R., Li, G., Zheng, F., & Tan, D. (2019). Humic Acid Fertilizer Improved Soil Properties and Soil Microbial Diversity of Continuous Cropping Peanut: A Three-Year Experiment. Scientific Reports, 9(1), 1–9. https://doi.org/10.1038/s41598-019-48620-4

Lindh, M. V., Lefébure, R., Degerman, R., Lundin, D., Andersson, A., & Pinhassi, J. (2015). Consequences of increased terrestrial dissolved organic matter and temperature on bacterioplankton community composition during a Baltic Sea mesocosm experiment. Ambio, 44, 402–412. https://doi.org/10.1007/s13280-015-0659-3

Littman, R. A., Bourne, D. G., & Willis, B. L. (2010). Responses of coral-associated bacterial communities to heat stress differ with Symbiodinium type on the same coral host. Molecular Ecology, 19(9), 1978–1990. https://doi.org/10.1111/j.1365-294X.2010.04620.x

Louvado, A., Cleary, D. F. R., Pereira, L. F., Coelho, F. J. R. C., Pousão-Ferreira, P., Ozório, R. O. A., & Gomes, N. C. M. (2021). Humic substances modulate fish bacterial communities in a marine recirculating aquaculture system. Aquaculture, 544, 737121. https://doi.org/10.1016/j.aquaculture.2021.737121

MacCarthy, P. (2001). The principles of humic substances. Soil Science, 166(11), 738–751. https://doi.org/10.1097/00010694-200111000-00003

MacNeil, M. A., Mellin, C., Matthews, S., Wolff, N. H., McClanahan, T. R., Devlin, M., Drovandi, C., Mengersen, K., & Graham, N. A. J. (2019). Water quality mediates resilience on the Great Barrier Reef. Nature Ecology and Evolution, 3(4), 620–627. https://doi.org/10.1038/s41559-019-0832-3

Mahmoud, H. M., & Kalendar, A. A. (2016). Coral-associated Actinobacteria: Diversity, abundance, and biotechnological potentials. Frontiers in Microbiology, 7, 204. https://doi.org/10.3389/fmicb.2016.00204

Manullang, C., Millyaningrum, I. H., Iguchi, A., Miyagi, A., Tanaka, Y., Nojiri, Y., & Sakai, K. (2020). Responses of branching reef corals Acropora digitifera and Montipora digitata to elevated temperature and pCO2. PeerJ, 8, e10562. https://doi.org/10.7717/peerj.10562

McDevitt-Irwin, J. M., Baum, J. K., Garren, M., & Vega Thurber, R. L. (2017). Responses of coralassociated bacterial communities to local and global stressors. Frontiers in Marine Science, 4, 1–16. https://doi.org/10.3389/fmars.2017.00262

Moberg, F., and Folke, C. (1999). Ecological goods and services of coral reef ecosystems. Ecol. Econ. 29, 215–233.

Mohamed, N. M., Saito, K., Tal, Y., & Hill, R. T. (2010). Diversity of aerobic and anaerobic ammonia-oxidizing bacteria in marine sponges. ISME Journal, 4(1), 38–48. https://doi.org/10.1038/ismej.2009.84

Montefalcone, M., Morri, C., & Bianchi, C. N. (2020). Influence of Local Pressures on Maldivian Coral Reef Resilience Following Repeated Bleaching Events, and Recovery Perspectives. Frontiers in Marine Science, 7, 1–14. https://doi.org/10.3389/fmars.2020.00587

Müller, R., Wiencke, C., Bischof, K., & Krock, B. (2009). Zoospores of three arctic laminariales under different UV radiation and temperature conditions: Exceptional spectral absorbance properties and lack of phlorotannin induction. Photochemistry and Photobiology, 85(4), 970–977. https://doi.org/10.1111/j.1751-1097.2008.00515.x

Mumby, P. J., Dahlgren, C. P., Harborne, A. R., Kappel, C. V., Micheli, F., Brumbaugh, D. R., Holmes, K. E., Mendes, J. M., Broad, K., Sanchirico, J. N., Buch, K., Box, S., Stoffle, R. W., & Gill, A. B. (2006). Fishing, trophic cascades, and the process of grazing on coral reefs. Science, 311(5757), 98–101. https://doi.org/10.1126/science.1121129

Nardi, S., Schiavon, M., & Francioso, O. (2021). Chemical structure and biological activity of humic substances define their role as plant growth promoters. Molecules, 26(8), 2256. https://doi.org/10.3390/molecules26082256

Otte, J. M., Blackwell, N., Soos, V., Rughöft, S., Maisch, M., Kappler, A., Kleindienst, S., & Schmidt, C. (2018). Sterilization impacts on marine sediment-Are we able to inactivate microorganisms in environmental samples? FEMS Microbiology Ecology, 94(12), 1–14. https://doi.org/10.1093/femsec/fiy189

Park, H. J., Lee, Y. M., Do, H., Lee, J. H., Kim, E., Lee, H., & Kim, D. (2021). Involvement of laccase-like enzymes in humic substance degradation by diverse polar soil bacteria. Folia Microbiologica, 66, 331–340. https://doi.org/10.1007/s12223-020-00847-9

Petersen, C. R., Jovanovic, N. Z., Le Maitre, D. C., & Grenfell, M. C. (2017). Effects of land use change on streamflow and stream water quality of a coastal catchment. Water SA, 43(1), 139–152. https://doi.org/10.4314/wsa.v43i1.16

Posadas, N., Baquiran, J. I. P., Nada, M. A. L., Kelly, M., & Conaco, C. (2022). Microbiome diversity and host immune functions influence survivorship of sponge holobionts under future ocean conditions. ISME Journal, 16(1), 58–67. https://doi.org/10.1038/s41396-021-01050-5

Ramaprasad, E. V. V., Mahidhara, G., Sasikala, C., & Ramana, C. V. (2018). Rhodococcus electrodiphilus sp. Nov., a marine electro active actinobacterium isolated from coral reef. International Journal of Systematic and Evolutionary Microbiology, 68(8), 2644–2649. https://doi.org/10.1099/ijsem.0.002895

Rautenberger, R., Wiencke, C., & Bischof, K. (2013). Acclimation to UV radiation and antioxidative defence in the endemic antarctic brown macroalga Desmarestia anceps along a depth gradient. Polar Biology, 36(12), 1779–1789. https://doi.org/10.1007/s00300-013-1397-2

Richardson, L.L., Ledrew, E.F., 2006. Remote sensing of aquatic coastal ecosystem processes. Sci. Manag. Appl. 9.

Roberts, C. M., McClean, C. J., Veron, J. E. N., Hawkins, J. P., Allen, G. R., McAllister, D. E., Mittermeier, C. G., Schueler, F. W., Spalding, M., Wells, F., Vynne, C., & Werner, T. B. (2002). Marine biodiversity hotspots and conservation priorities for tropical reefs. Science, 295(5558), 1280–1284. https://doi.org/10.1126/science.1067728

Rocha, R. J. M., Bontas, B., Cartaxana, P., Leal, M. C., Ferreira, J. M., Rosa, R., Serôdio, J., & Calado, R. (2015). Development of a Standardized Modular System for Experimental Coral Culture. Journal of the World Aquaculture Society, 46(3), 235–251. https://doi.org/10.1111/jwas.12186

Rocha, R. J. M., Rodrigues, A. C. M., Campos, D., Cícero, L. H., Costa, A. P. L., Silva, D. A. M., … & Silva, A. P. (2020). Do microplastics affect the zoanthid Zoanthus sociatus? Science of The Total Environment, 713, 136659.

Rocker, D., Kisand, V., Scholz-Böttcher, B., Kneib, T., Lemke, A., Rullkötter, J., & Simon, M. (2012). Differential decomposition of humic acids by marine and estuarine bacterial communities at varying salinities. Biogeochemistry, 111(1–3), 31–346. https://doi.org/10.1007/s10533-011-9653-4

Rong, J. C., Ji, B. W., Zheng, N., Sun, Z. Z., Li, Y. S., & Xie, B. Bin. (2021). Genomic insights into antioxidant activities of Pyruvatibacter mobilis CGMCC 1.15125T, a pyruvate-requiring bacterium isolated from the marine microalgae culture. Marine Genomics, 55, 100791. https://doi.org/10.1016/j.margen.2020.100791

Rosado, P. M., Leite, D. C. A., Duarte, G. A. S., Chaloub, R. M., Jospin, G., Nunes da Rocha, U. P. Saraiva, J., Dini-Andreote, F., Eisen, J. A., Bourne, D. G., & Peixoto, R. S. (2019). Marine probiotics: increasing coral resistance to bleaching through microbiome manipulation. ISME Journal, 13(4), 921–936. https://doi.org/10.1038/s41396-018-0323-6

Roth, M. S. (2014). The engine of the reef: Photobiology of the coral-algal symbiosis. Frontiers in Microbiology, 5, 1–22. https://doi.org/10.3389/fmicb.2014.00422

Sharpless, C. M., Aeschbacher, M., Page, S. E., Wenk, J., Sander, M., & McNeill, K. (2014). Photooxidation-induced changes in optical, electrochemical, and photochemical properties of humic substances. Environmental Science and Technology, 48(5), 2688–2696. https://doi.org/10.1021/es403925g

Shore, A., Sims, J. A., Grimes, M., Howe-Kerr, L. I., Grupstra, C. G. B., Doyle, S. M., Stadler, L., Sylvan, J. B., Shamberger, K. E. F., Davies, S. W., Santiago-Vázquez, L. Z., & Correa, A. M. S. (2021). On a Reef Far, Far Away: Anthropogenic Impacts Following Extreme Storms Affect Sponge Health and Bacterial Communities. Frontiers in Marine Science, 8, 693430. https://doi.org/10.3389/fmars.2021.608036

Spaccini, R., Mbagwu, J. S. C., Conte, P., & Piccolo, A. (2006). Changes of humic substances characteristics from forested to cultivated soils in Ethiopia. Geoderma, 132(1-2), 9–19. https://doi.org/10.1016/j.geoderma.2005.04.015

Stoms, D. M., Davis, F. W., Andelman, S. J., Carr, M. H., Gaines, S. D., Halpern, B. S., … & Warner, R. R. (2005). Integrated coastal reserve planning: making the land–sea connection. Frontiers in Ecology and the Environment, 3(8), 429–436.

Stuij, T. M., Cleary, D. F. R., Rocha, R. J. M., Polonia, A. R. M., Silva, D. A. M., Frommlet, J. C., Louvado, A., Huang, Y. M., van der Windt, N., de Voogd, N. J., & Gomes, N. C. M. (2023). Development and validation of an experimental life support system to study the impact of ultraviolet B radiation and temperature on coral reef microbial communities. BioRxiv, 2023.03.01.530425. https://doi.org/10.1101/2023.03.01.530425

Sun, F., Yang, H., Zhang, X., Tan, F., & Shi, Q. (2022). Response characteristics of bacterial communities in multiple coral genera at the early stages of coral bleaching during El Niño. Ecological Indicators, 144, 109569. https://doi.org/10.1016/j.ecolind.2022.109569

Sweet, M., Ramsey, A., & Bulling, M. (2017). Designer reefs and coral probiotics: great concepts but are they good practice? Biodiversity, 18(1), 19–22.

Teichberg, M., Wild, C., Bednarz, V. N., Kegler, H. F., Lukman, M., Gärdes, A. A., Heiden, J. P., Weiand, L., Abu, N., Nasir, A., Miñarro, S., Ferse, S. C. A., Reuter, H., & Plass-Johnson, J. G. (2018). Spatio-temporal patterns in coral reef communities of the Spermonde Archipelago, 2012-2014, I: Comprehensive reef monitoring of water and benthic indicators reflect changes in reef health. Frontiers in Marine Science, 5, 1–18. https://doi.org/10.3389/fmars.2018.00033

Torti, A., Lever, M. A., & Jørgensen, B. B. (2015). Origin, dynamics, and implications of extracellular DNA pools in marine sediments. Marine Genomics, 24, 185–196. https://doi.org/10.1016/j.margen.2015.08.007

Travesso, M., Missionário, M., Cruz, S., Calado, R., & Madeira, D. (2023). Combined effect of marine heatwaves and light intensity on the cellular stress response and photophysiology of the leather coral Sarcophyton cf. glaucum. Science of the Total Environment, 861, 160460. https://doi.org/10.1016/j.scitotenv.2022.160460

Trench, R. K. (1979). The cell biology of plant-animal symbiosis. Annual Review of Plant Physiology, 30(1), 485–531.

Voolstra, C. R., & Ziegler, M. (2020). Adapting with Microbial Help: Microbiome Flexibility Facilitates Rapid Responses to Environmental Change. BioEssays, 42(7), 1–9. https://doi.org/10.1002/bies.202000004

Wessels, W., Sprungala, S., Watson, S. A., Miller, D. J., & Bourne, D. G. (2017). The microbiome of the octocoral Lobophytum pauciflorum: Minor differences between sexes and resilience to short-term stress. FEMS Microbiology Ecology, 93(5), fix013. https://doi.org/10.1093/femsec/fix013

Weyrich, L. S., Farrer, A. G., Eisenhofer, R., Arriola, L. A., Young, J., Selway, C. A., Handsley-Davis, M., Adler, C. J., Breen, J., & Cooper, A. (2019). Laboratory contamination over time during low-biomass sample analysis. Molecular Ecology Resources, 19(4), 982–996. https://doi.org/10.1111/1755-0998.13011

Wyatt, A. S. J., Leichter, J. J., Toth, L. T., Miyajima, T., Aronson, R. B., & Nagata, T. (2020). Heat accumulation on coral reefs mitigated by internal waves. Nature Geoscience, 13(1), 28–34. https://doi.org/10.1038/s41561-019-0486-4

Ying, J., Collins, M., Cai, W., Timmermann, A., Huang, P., Chen, D., & Stein, K. (2022). Emergence of climate change in the tropical Pacific. Nature Climate Change, 12(4), 356–364. https://doi.org/10.1038/s41558-022-01301-z

Ziegler, M., Grupstra, C. G. B., Barreto, M. M., Eaton, M., BaOmar, J., Zubier, K., Al-Sofyani, A., Turki, A. J., Ormond, R., & Voolstra, C. R. (2019). Coral bacterial community structure responds to environmental change in a host-specific manner. Nature Communications, 10(1), 3092. https://doi.org/10.1038/s41467-019-10969-5

Zweifler, A., O’Leary, M., Morgan, K., & Browne, N. K. (2021). Turbid Coral Reefs: Past, Present and Future—A Review. Diversity, 13(6), 251. https://doi.org/10.3390/d13060251

