## supplementary Fig. 1 for "Humic substances mitigate adverse effects of elevated temperature with potentially critical repercussions for coral reef resilience"

Postal address:
Daniel Cleary & Newton Gomes
Centre for Environmental and Marine Studies (CESAM)
Second floor, building 2
Campus Universitário Santiago,
3810-193 Aveiro, Portugal

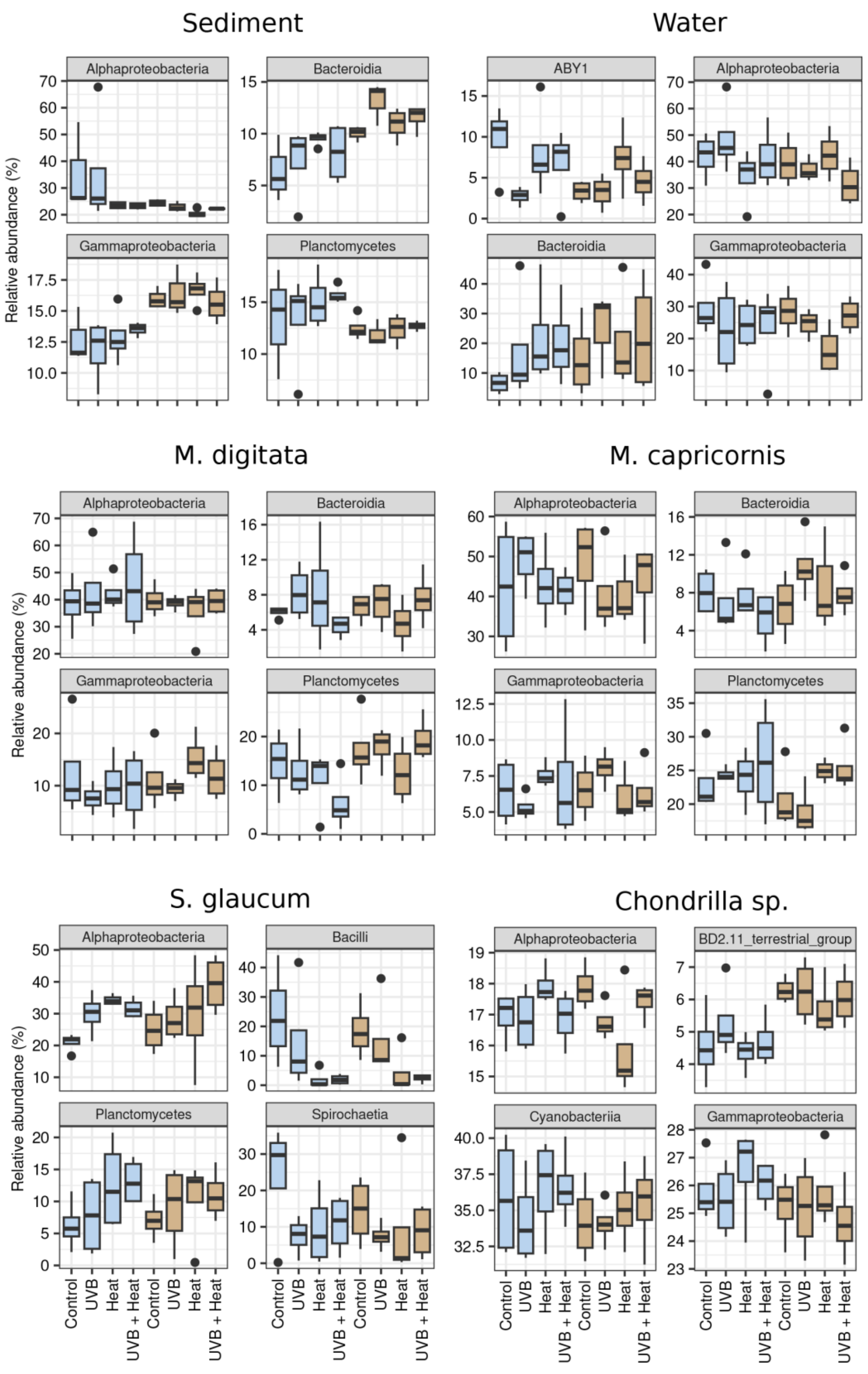


**Supplementary Fig. 1** Boxplots of the relative abundance of the four most abundant classes in sediment, water, *M. digitata, M. capricornis, S. glaucum* and *Chondrilla sp.* under the independent and combined effects of UVB, Heat and HS supplementation. Relative abundances are grouped per treatment within each biotope. Boxplots of treatments without HS supplementation are depicted in blue, boxplots of treatments with HS supplementation are depicted in brown.
